# On the mechanism of K^+^ transport through the inter-subunit tunnel of KdpFABC

**DOI:** 10.1101/2025.03.04.641403

**Authors:** Hridya Valia Madapally, Adel Hussein, Martin Wazar Eriksen, Bjørn Panyella Pedersen, David L. Stokes, Himanshu Khandelia

## Abstract

KdpFABC is an ATP-dependent membrane complex that enables prokaryotes to maintain potassium homeostasis under potassium-limited conditions. It features a unique hybrid mechanism combining a channel-like selectivity filter in KdpA with the ATP-driven transport functionality of KdpB. A key unresolved question is whether K^+^ ions translocate through the inter-subunit tunnel as a queue of ions or within a hydrated environment. Using molecular dynamics (MD) simulations, metadynamics, anomalous X-ray scattering, and biochemical assays, we demonstrate that the tunnel is predominantly occupied by water molecules rather than multiple K^+^ ions. Our results identify only one stable intermediate binding site for K^+^ within the tunnel, apart from the canonical sites in KdpA and KdpB. Free energy calculations reveal a substantial barrier (∼22 kcal/mol) at the KdpA-KdpB interface, making spontaneous K^+^ translocation unlikely. Furthermore, mutagenesis and functional assays confirm previous findings that Phe232 at this interface plays a key role in coupling ATP hydrolysis to K^+^ transport. These findings challenge previous models proposing a continuous wire of K^+^ ions through the tunnel and suggest the existence of an as-yet unidentified intermediate state or mechanistic detail that facilitates K^+^ movement into KdpB.

## INTRODUCTION

In potassium-deficient environments, prokaryotes rely on the ATP-dependent membrane protein KdpFABC to transport potassium ions against a large transmembrane electrochemical gradient to maintain intracellular K^+^ homeostasis. KdpFABC is a unique hybrid of a channel and a pump comprising four protein subunits^1^. Subunit KdpA descends from a superfamily of K^+^ transporters^2^ and consists of a K^+^ channel-like selectivity filter that distinguishes K^+^ from other ions. KdpA has 10 helices organized into four tandem repeats of the characteristic MPM motif that characterizes K^+^ channels. The KdpB subunit, evolutionarily and structurally a P-type ATPase pump^3^, uses the Post-Albers cycle to couple ATP hydrolysis to transport^4, 5^. KdpB consists of a transmembrane domain with seven helices harboring a canonical ion binding site. The cytoplasmic A, P, and N domains undergo extensive structural changes that are coupled to ATP binding, phosphorylation of a conserved aspartate, followed by its hydrolysis. KdpC is closely associated with KdpA and consists of a soluble periplasmic domain with a single transmembrane helix. The function of KdpC is unknown and has been speculated to influence substrate affinity. KdpF consists of a single transmembrane helix proposed to stabilize the complex. This subunit is not universally conserved and the *E. coli* protein remains functional without KdpF in lipid-containing environments or membranes.^6^ As a complex, KdpFABC features a ∼40 Å long tunnel extending from KdpA’s selectivity filter to KdpB’s substrate binding site, which is one of its most intriguing structural characteristics.^1^

KdpFABC was initially identified as an ATP-dependent pump due to the P-type ATPase-like properties of KdpB, which could be phosphorylated, contained conserved pump-specific motifs, and exhibited vanadate-sensitive ATPase activity.^7, 8^ Later studies revealed that mutations in KdpA significantly reduced transport efficiency, indicating that KdpA played a critical role in the active transport of K^+^.^9, 10^ This led to the proposition that KdpA regulated selectivity and transport while KdpB provided the energy for uphill K^+^ import via ATP hydrolysis, while ensuring that transport and ATP hydrolysis remained coupled.

The coupling helix model^1^ was initially proposed as a mechanism in which transport and ATP hydrolysis occur in spatially distinct regions of KdpFABC. It suggested that the tunnel between the KdpA and KdpB subunits is filled with water molecules, forming a “water wire” that mediates proton transfer between KdpA and KdpB. K^+^ binding at the S3 site of the KdpA selectivity filter triggers proton movement towards KdpB which, upon reaching the canonical binding site in KdpB, initiates ATP hydrolysis. The conformational change induced by ATP hydrolysis would pull on an intramembrane gate in KdpA via a salt-bridge network linking the cytoplasmic loops of KdpA and KdpB. The consequent opening of this KdpA gate would allow K^+^ to move into the cytoplasm. However, cryo-EM structures later revealed that KdpA remained structurally invariant during various stages of the Post-Albers cycle.^11, 12^ Additionally, the salt bridge between KdpA and KdpB was absent in the E2 state of the protein, and mutations in the salt-bridge residues did not affect transport^12^. The coupling helix model was thus not the answer to the transport mechanism.

Instead, the cryo-EM structures led to the proposal of a new tunnel translocation model.^11^ In this model, K^+^ translocates from the selectivity filter of KdpA to the canonical binding site in KdpB during the E1-ATP state via the aforementioned intersubunit tunnel. K^+^ binding at the KdpB site would trigger transfer of phosphate from ATP to the conserved aspartate in the P-domain. The subsequent transition from E1-P to E2-P pinches off the tunnel while moving the K^+^ ion to a nearby low-affinity site in the transmembrane binding site of KdpB, from which it is finally released into the cytoplasm. The tunnel translocation model raises an important question about tunnel occupancy. In one scenario, the tunnel is constitutively filled with K^+^ ions up to the KdpA-KdpB interface, and the binding of an additional K^+^ with accompanying ATP hydrolysis propels the final K^+^ ion at the pump-channel interface into the canonical KdpB binding site^13^. In an alternative scenario, the tunnel is mainly occupied by water (not necessarily in a single file), allowing K^+^ to move through a hydrated tunnel, also in conjunction with ATP hydrolysis^12^. Our study is designed to distinguish between these two scenarios.

Cryo-EM structures solved by two different groups identified multiple densities within the inter-subunit tunnel.^12, 13^ However, it remains unclear whether these densities represent K^+^ or water. Sweet and co-workers^12^ attributed most of these densities to water molecules due to a general lack of compensating charges in a region of low dielectric. In contrast, Silberberg and co-workers^13^ assigned the densities to K^+^ ions based on effects of Rb^+^ substitution and MD simulations. Furthermore, these simulations showed a K^+^ ion diffusing spontaneously from the pump-channel interface to the binding site in KdpB. However, the diffusion was observed in a rather short simulation (within 200 ps) in which the peptide backbone was constrained, and the tunnel was prepopulated by multiple K^+^ ions, which can lead to unrealistic electrostatic repulsion. Accurate free energy calculations to quantify the energetics of the transfer were not performed. It remains uncertain whether such a transfer will occur if the tunnel is not prepopulated by K^+^ ions: a state which may or may not be realistic. Furthermore, the possibility of such a transfer will depend on the energetics of the coupling of ATP hydrolysis to the transport of K^+^ across the KdpA-KdpB interface.

Here, we investigate the contents of the tunnel and the energetics of K^+^ transport through the KdpFABC complex using molecular dynamics (MD) simulations, free energy calculations, anomalous X-ray scattering and biochemical analysis of relevant mutants. MD simulations indicate that there is only one favorable binding site for K^+^ ion in the tunnel region in KdpA. Numerous water molecules spontaneously occupy the tunnel, but the water wire is not continuous due to the presence of a hydrophobic barrier at the pump-channel interface. The same hydrophobic region also creates a high free energy barrier for movement of K^+^ through the tunnel from the S3 site to the canonical binding site in KdpB in the E1-ATP state. We did not investigate the E2 conformation because the tunnel is known to be partially collapsed in this state.^11^ Supporting these simulations, anomalous X-ray scattering data detected K^+^ in the selectivity filter but not in the tunnel nor at the canonical KdpB binding sites. As in previous work^13^, both MD simulations and functional assays indicate that Phe232 at the subunit interface plays an important role in K^+^ translocation through the tunnel. For clarity, we will refer to the region comprising both the selectivity filter and the inter-subunit tunnel as the “translocation passage” whereas “tunnel” solely refers to the intramembrane cavity and excludes the selectivity filter.

## METHODS

### Simulation system

The initial configuration of the protein was derived from the PDB ID: 7LC3 (E1-ATP). The PDB ID 7LC3 corresponds to E1-ATP state of the protein. The E1-ATP-F232A state was obtained by F232A mutation in KdpB subunit of 7LC3. The protein coordinates along with the ATP/Pi molecule and Mg^2+^ ion were extracted and mutations were reversed to create the wildtype protein. K^+^ ions were placed at different positions in the selectivity filter and tunnel for a series of simulations as described later. The protein orientation in the membrane was identified using the OPM server^14^. CHARMM-GUI^15-17^ was used to insert the protein into a symmetric bilayer consisting of 98 POPG lipids, 49 Cardiolipins and 344 POPE lipids. Since POPE lipids are the most abundant, the lipid density near the interface of KdpA and KdpB was modelled as a POPE lipid as in the deposited structure. The system was solvated with TIP3P^18^ water molecules and an ion concentration of 150 mM was maintained using K^+^ and Cl^-^ ions. The system was energy minimized using the steepest descent method. Three short NVT and NPT equilibrations of equal intervals were performed for a total of 15 ns, while gradually releasing the position restraints on the heavy atoms of the protein and membrane. In cases where K^+^ ions are present in the translocation passage; the system was further equilibrated for 100 ns while position restraints were applied only on the K^+^ ions placed at different locations in the translocation passage. The resulting structure was simulated for 1 μs without any restraints at a temperature of 310 K and 1 bar using the Nose-Hoover thermostat^19, 20^ and Parinello-Rahman barostat^21^, respectively. Multiple replicas of each system were simulated. Electrostatic interactions were treated using the PME^22^ method with a cut-off range of 1.2 nm and the van der Waals interaction was switched off between 1.0 and 1.2 nm using the force-switch function. The CHARMM36 forcefield^23-25^ was used for the components in the system and all the simulations were performed using GROMACS-2021.4^26^.

### Metadynamics simulations

Multiple walker well-tempered metadynamics simulations^27^ were performed using GROMACS patched with PLUMED^28, 29^. The binding site in KdpB (hereafter referred to as B site) was defined as the center of the mass of the atoms 261CYS-O, 263ILE-O and 583ASP-OD1. The projection of the distance between the K^+^ ion and the above-mentioned center of mass in the KdpB binding site on the *x*-axis was the collective variable (CV) employed in the metadynamics simulations. We have used projection rather than the distance since it allows us to identify the exact position of the ion along the tunnel. A higher value of the CV indicates that the ion is further away from the binding site, B while a smaller value indicates that the ion is close to B. CV=−0.2 indicates that ion is bound at B. Further lower negative values of the CV indicates that the ion has crossed the B site in KdpB. Five replicas were simulated with different initial positions of the K^+^ ion in the translocation passage. The initial placements of the ions were at S3, S4, I1, B sites and at the subunit interface. A hill height of 2.0 kJ/mol with a sigma of 0.015 was deposited every 1 ps with a bias factor of 20. Walls were defined to avoid sampling of the K^+^ outside the tunnel region.

### Anomalous X-ray Scattering

For crystallization, protein was produced and purified as described previously^1^. Briefly, KdpFABC carrying the Q116R mutation in KdpA was expressed from the pSD107 plasmid using K^+^ limitation to activate its native promoter in E. coli strain TK2498 [F-thi rha lacZ nagA trkA405 trkD1 Δ(kdpFABC)5 Δ(ompT)]. After cell disruption using an Emulsiflex C3 homogenizer (Avestin), the membrane fraction was solubilized with n-decyl-*β*-maltoside (DM, 0.24 g/ gram of membrane pellet) and the His-tagged KdpFABC complex was isolated using a Ni-NTA HisTrap column (GE Healthcare). The complex was then purified by size exclusion chromatography using a Superdex 200 increase 10/300 column (GE Healthcare) equilibrated with a buffer containing 100 mM KCl, 25 mM Tris pH 7.5, 10% glycerol, 1% octyl glucoside and 0.5 mg/ml dimyristoylphosphatidylcholine. Peak fractions were combined, concentrated to 12 mg/ml and then set up for crystallization using hanging drop vapor diffusion against buffers containing 15-25% PEG3350, 5 mM AMP-PCP, 0.5-0.8 M NaCl, 0.05 M sodium citrate pH 5.5. Crystals were frozen in liquid nitrogen and shipped to MAX IV synchrotron (Lund, Sweden) for data collection at the BioMax beamline. X-ray diffraction data were collected at 7 keV (1.77 A wavelength) using a helical scan with 0.1° intervals for a total of 360°. Data were processed with the automated pipeline at BioMax employing the staraniso algorithm to account for anisotropy (globalphasing.org). Phasing and generation of the anomalous map were done with PHENIX^30^. First, the previously deposited X-ray structure corresponding to this crystal form (5MRW) was used to phase the data. A single round of refinement was done to ensure that the model was consistent with the data. Finally, structure factors, including those for the anomalous signal, were generated with phenix.find_peaks_holes. Although data extended to 3.1 Å resolution (Supplementary Table. 1), it was truncated to 3.3 Å for this last step, which was observed to produce maximal anomalous signal in the selectivity filter.

### Activity measurements

For functional assays, wild-type (WT) and mutant KdpFABC complexes were expressed as previously described^31^ from a pBAD vector in bacterial strain TK2281(ΔkdpFABCDE81 thi rha lacZ nagA trkA405 trkD1). The TK2281 strain is notable in lacking both kdpFABC operon and the kdpDE operon, the latter of which is responsible both for inducing expression of the endogenous complex under physiological conditions and for post-translational inhibition via phosphorylation of Ser162 on KdpB^32^. By using a pBAD vector for heterologous expression in TK2281, both active and inactive protein constructs can be produced free from the inhibitory effects of Ser162 phosphorylation. Expression and purification protocols are described by Hussein et al.^31^. In brief, after solubilization of cell membranes by DM and isolation by Ni-NTA affinity chromatography, as described above, final purification was performed by size-exclusion chromatography using a Superdex 200 column and a buffer containing 100 mM NaCl, 25 mM Tris (pH 7.5), 10% glycerol, 1 mM TCEP, and 0.15% DM.

For measurements of ATPase activity, a coupled enzyme assay was performed as described previously^32^ in a buffer containing 75 mM TES pH 7, 150 mM KCl, 7.5 mM MgCl_2_, 9.6 U/mL lactate dehydrogenase, 9.6 U/mL pyruvate kinase, 2.4 mM ATP, 0.5 mM phosphoenol-pyruvate, 0.36 mM NADH and 0.15% DM. For measurement of K^+^ transport, protein was reconstituted as previously described^31^ into lipid vesicles containing a 4:1 weight ratio of 1-palmitoyl-2-oleoyl-sn-glycero-3-phosphocholine and 1,2-dioleoyl-sn-glycero-3-phosphate (Avanti Polar Lipids, Alabaster AL). These proteoliposomes, containing a lipid-to-protein weight ratio of 10:1, were adsorbed to gold-plated sensors for solid-supported membrane electrophysiology (SSME) measurements of charge transfer using the SURFE^2^RN1 (Nanion Technologies, Livingston NJ).

## RESULTS

### K^+^ ions in the translocation passage

To explore whether the cryo-EM densities in the tunnel^12, 13^ correspond to K^+^ ions or water molecules, we launched simulations of the E1-ATP state, starting with a different number of K^+^ ions in the translocation passage. In the following, we will describe the fate of K^+^ ions along the entire translocation passage as the initial number of K^+^ ions in the tunnel are varied.

1 K^+^ in the translocation passage (single simulation) – We started by placing a K^+^ ion at the S3 site because anomalous signal from the initial X-ray structure showed the presence of K^+^ at that location (this result is confirmed by new data shown below). During the simulation, K^+^ remains at the S3 site for the entire 1 μs (Figure. 1a,b), and is coordinated by carbonyl oxygen atoms of N112, T113, T230, S343, C344, N466 and N467 in agreement with the cryo-EM structures (Figure S1.(a),(b)). S3 is thus a stable binding site for K^+^ as previously deduced from structural data.

**Figure 1:**
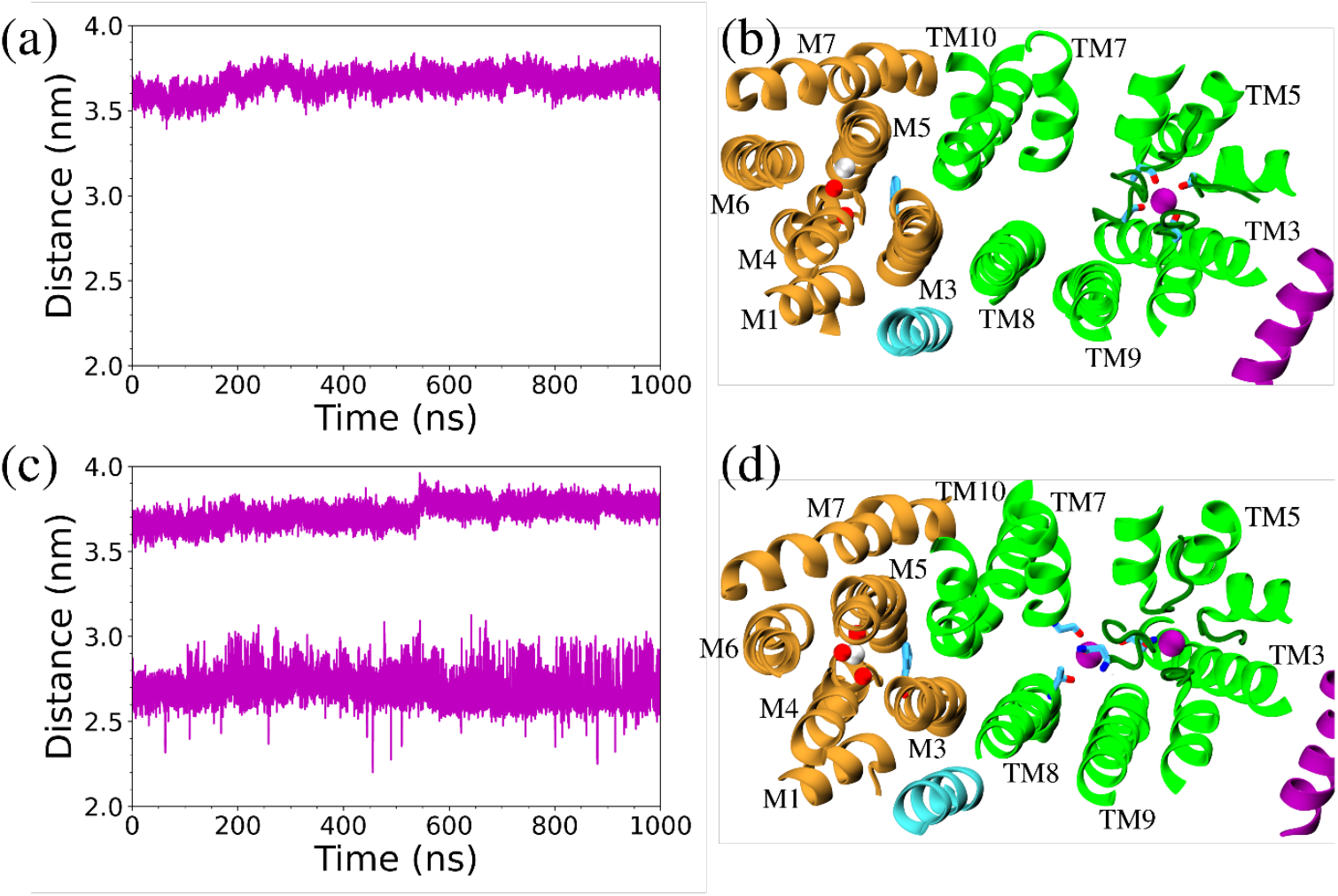
(a) and (c) Distance between the K^+^ ions and the center of mass of the KdpB ion-binding site in systems with 1 and 2 K^+^ ions placed initially in the translocation passage. In (b) and (d), the red spheres represent the oxygens atoms that form the KdpB binding site and the white sphere represents the center of mass of those oxygen atoms. The purple spheres represent K^+^ ions at S3 and I1. KdpC, KdpB, KdpA, and KdpF are shown in purple, brown, green, and cyan respectively. Selectivity filter loops are depicted in dark green color.

2 K^+^ in the translocation passage (single simulation) – We placed the first K^+^ at the S3 site and the second K^+^ in the tunnel at an arbitrary position between the S3 and the interface of KdpA and KdpB. During the simulation, the ion at S3 remained in its initial position, whereas the second ion quickly moved to an intermediate location between the S3 site and the interface (Figure. 1c,d). We refer to this new location as site I1, where the K^+^ was coordinated by oxygen atoms of the residues G369, S378, T424 and N465 of KdpA (Figure S1.(c),(d)). The ion appeared to be stably bound at I1, though with somewhat larger deviations relative to S3. The two ions remain at their respective sites and do not exchange their positions during the simulation.

3 K^+^ in the translocation passage (four simulations) – In this setup, the first and second K+ ions were positioned at the S3 and I1 sites as they were identified to be stable sites from the previous simulations. In two of the simulations we placed the third ion between I1 and the subunit interface, whereas in other two sets, the third ion was placed between S3 and I1 (see Figure S2). In three of the four simulations, one K^+^ ion escapes to the solvent through the KdpA selectivity filter (Supplementary Movie-1). The K^+^ ion closest to the mouth of the selectivity filter (at S3) is the ion that escapes into the bulk; this results in instantaneous settlement of another K^+^ ion from the translocation passage into the empty S3 site. However, in one simulation (Simulation #3 in Figure. 2a), three K^+^ ions remained in the translocation passage for 1 μs, where the third ion settled into a new position close to the S4 site of the selectivity filter. The ion at the S4 site was coordinated by oxygen atoms of the residues A342, S343, G374, S378, and N466 of KdpA (Figure S1.(e),(f)). The ions at I1 and S4 occasionally exchanged their positions. These simulations suggest that aside from the K+ at the KdpB binding site, only one K+ ion can occupy the I1 site within the tunnel, while the remaining K+ ions are accommodated within the selectivity filter.

**Figure 2:**
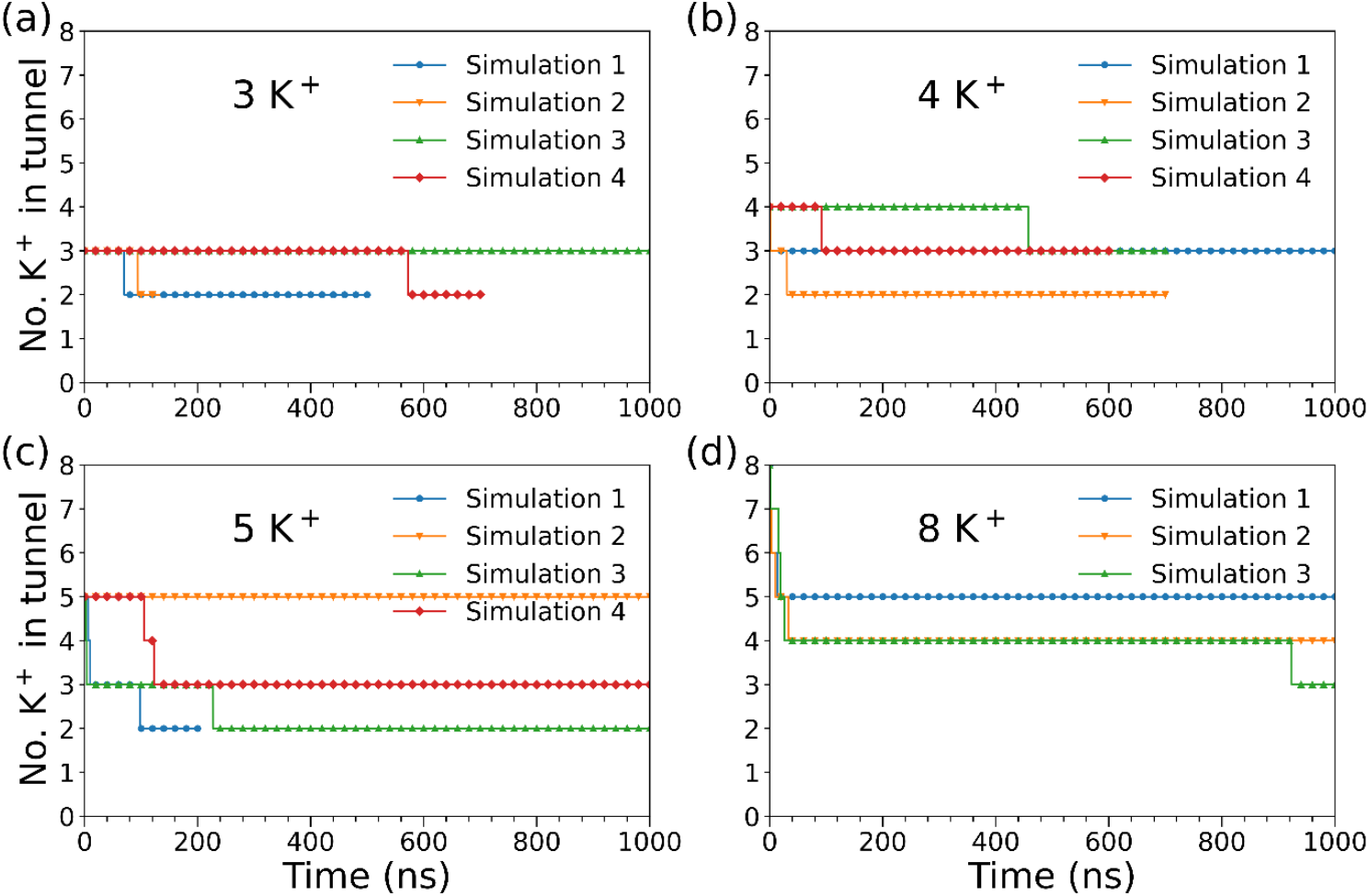
Number of K^+^ ions that remain in the translocation passage in simulations starting with different initial number of K^+^ ions.

4 K^+^ in the translocation passage (four simulations) – We simulated four systems with four K^+^ ions in the translocation passage. In two simulations, the ions were initially positioned at the S2, S3, I1 sites, and a location between I1 and the interface. In the other two simulations, the ions were placed at S3, S4, I1, and the same position between I1 and the interface. During these simulations, one or two K+ ions escaped through the selectivity filter, leaving only two or three ions in the translocation passage (Figure 2b). When three ions remained in the translocation passage, one was located in the tunnel (I1), while the other two occupied the S3 and S4 sites within the selectivity filter. When two ions remained in the translocation passage, they occupied S3 and I1 site. These data again indicate that only one K+ ion (at I1 site) can occupy the tunnel in KdpA region, while the remaining K+ ions are positioned in the selectivity filter.

5 K^+^ in the translocation passage (four simulations)– We simulated four systems with 5 K^+^ ions in the translocation passage. A K^+^ ion was placed at S3, S4, and I1 site each and rest of the two ions were arbitrarily placed in the tunnel within the KdpA section of the translocation passage in all the systems. In three systems, two or more K+ ions escaped via the selectivity filter (Figure 2c) thus ensuring that at most 3 K^+^ ions stayed in the translocation passage (at S3, S4 and I1 in Simulation #4 and at S3 and I1 in Simulation #1 and #3). However, in one of the simulations (Simulation #2), all five ions remained in the passage. A closer analysis revealed that in this simulation, chloride ions from the solvent entered the tunnel, stabilizing the high concentration of K+ ions (Figure S3 and Supplementary Movie 2). The presence of these chloride ions is an artifact caused by crowding the passage with too many K+ ions, which creates an energetically unfavorable state. Previous investigations of the Na^+^, K^+^ pump have shown that the binding pocket is unstable under such conditions, either expelling ions or attracting negatively charged ions from the bulk.^33^ These data suggest that it is highly unlikely for five K+ ions to simultaneously occupy the translocation passage.

8 K^+^ in the translocation passage (three simulations)– We simulated three systems with 8 K^+^ ions in the translocation passage. In one of the simulations, the ions were positioned based on the densities observed in PDB ID 7NNL (the K^+^ ions at the KdpB binding site and at the subunit interface in 7NNL were removed), which corresponds to the E1-ATP state of KdpB^13^. In the other two systems, a K^+^ ion was placed each at S3, S4 and I1 sites and the rest of the ions were placed in the KdpA section of the translocation passage. The simulations with eight K+ ions produced results similar to those with five K+ ions (Figure 2d). In one of the simulations, only three ions remained in the translocation passage while other ions escaped into the bulk. In the other two systems, the translocation passage attracted chloride ions from the solvent into the tunnel.

All the above simulations provide evidence against the presence of more than 3 K^+^ ions in the translocation passage, and more than 1 K^+^ ion within the tunnel itself, in the E1-ATP state. Most of the spherical densities observed in the cryo-EM structures are therefore likely to represent tightly co-ordinated water molecules (more on this later). Additionally, the failure of any K+ ion to translocate to the KdpB binding site, even with eight ions in the passage, points to the presence of a free energy barrier at the interface between the two subunits.

### Simulation of Phe232Ala (F232A) mutation with 8 K^+^ ions in the passage

Phe232 is a conserved residue of KdpB, and its mutation to either Ile or Ala has been reported to reduce ion transport efficiency^13^ and, in the case of the F232A mutant, to uncouple ATP hydrolysis from transport. The bulky Phe232 sidechain can create a constriction just before the binding site in KdpB and has been hypothesized to aid in ion coordination at B through direct cation-π interactions or CH-π interactions with L262 of KdpB^13^. To explore whether removing the constriction formed by Phe232 would reduce the barrier at the interface and allow easier passage of K^+^ ions from KdpA to KdpB along the tunnel, we conducted simulations with the F232A mutation with 8 K^+^ in the passage. However, during the 1 μs simulation, the ions did not spontaneously cross the interface to reach the KdpB binding site. Instead, several ions escaped through the selectivity filter (Figure S4) leaving only three ions in the translocation passage (at S3, S4 and I1 site). The simulation provides evidence that the free energy barrier at the interface is not relieved by F232A mutation in KdpB.

### Free energy of K^+^ translocation along the tunnel

To better understand the lack of K^+^ translocation events from KdpA to KdpB, we quantified the free energy barrier at the subunit interface. For this, we launched multiple-walker well-tempered metadynamics simulations to estimate the free energy required for K^+^ to move through the tunnel from the S3 site in KdpA to the binding site (B) in KdpB. The collective variable (CV) used was the projection of the distance between the K^+^ ion and the B site along the *x*-axis, which nearly aligns with the tunnel’s direction. We generated free energy profiles for the E1-ATP and E1-ATP-F232A states. The converged free energy values are shown in Figure 3a (see Figures S5-S8 for free energy convergence and additional calculation details). CV values at 3.45, 2.65, and - 0.20 nm correspond to the S3, I1, and B sites, respectively. The zero in the free energy profile is at I1 site. There are distinct minima at defined ion binding sites within the selectivity filter (*d.x* ≈ 3.45*nm*) and in KdpB (*d.x* ≈ −0.20*nm*). The minima at *d.x* ≈ 2.65*nm* corresponding to I1 site aligns with the findings from the simulations with multiple K^+^ ions in the passage, which demonstrated that K^+^ ion can bind only a single site in tunnel in KdpA, apart from the ions in the selectivity filter. The minima at *d.x* ≈ −0.2*nm* is at the canonical ion binding site in the KdpB subunit (see Figure. 3c).

**Figure 3:**
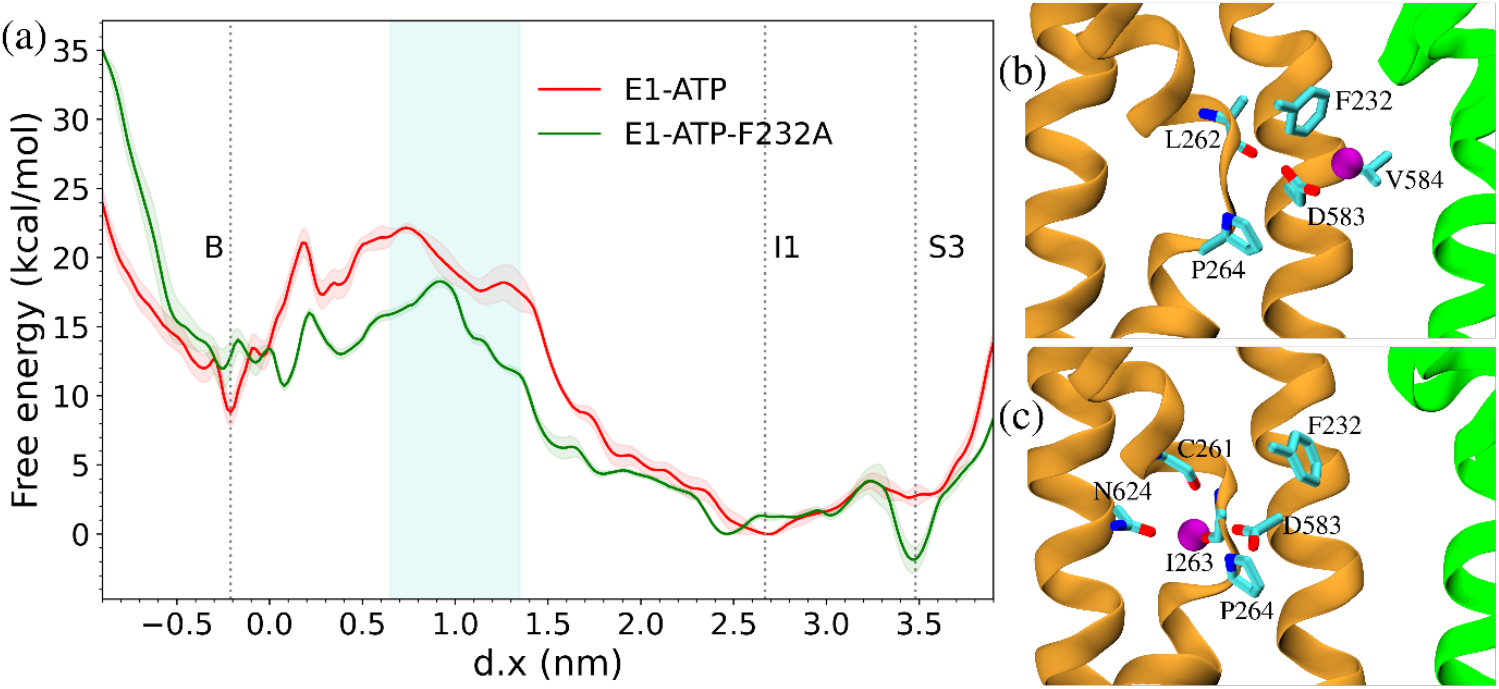
(a) Free energy of passage of K^+^ along the tunnel obtained using multiple walker well-tempered Metadynamics. *d.x* ≈ −0.20 *nm* corresponds to the ion binding site, B in KdpB, *d.x* ≈ 3.45 *nm* corresponds to the S3 site of KdpA and *d.x* ≈ 2.65 *nm* corresponds to the I1 site at KdpA. The I1 site has been chosen as the zero of the free energy. The light blue shaded region corresponds to the transition state region. (b) The transition state at *d.x* ≈ 0.60 *nm*. The K^+^ ion is at the interface and is yet to cross Phe232. (c) K^+^ ion at the B site coordinated by Asn624, Asp583, Cys261 and Ile263.

This high energy barrier likely arises due to the presence of a hydrophobic expanse at this region. The CV spanning from *d.x* ≈ 1.35 nm to *d.x* ≈ 0.65*nm* corresponds to a transition region corresponding to the subunit interface (light blue shaded region in Figure 3a). The K^+^ ion remains uncoordinated by oxygen from the side chain of the amino acids and water molecules in this region. Thus, the high barrier results from the desolvation of K^+^ at the hydrophobic interface within the tunnel between KdpA and KdpB. For instance, the barrier is Δ*G* = 22.17 *kcal*/*mol* for the E1-ATP system. A similarly high barrier is observed in the E1-ATP-F232A system.

### Hydration of the tunnel

Our calculations indicate that the translocation pathway is only able to accommodate 2-3 K^+^ ions with only one K^+^ occupying the tunnel at the I1 site. In contrast, we observed many water molecules populating the tunnel in the simulations (Figure S9). To quantify tunnel hydration, we first identified the tunnel region in each trajectory frame using the program HOLE^34^. Our procedure accounts for the fact that the position of the tunnel fluctuates between frames due to thermal motion. We then use a custom code to count the number of water molecules within the tunnel. Figure 4a shows the number of water molecules (data represents a running average of 20 frames) present in the tunnel for simulations with one K^+^ ion (at S3) and three K^+^ ions (at S3, S4, and I1) in the translocation passage. The average number of water molecules in the tunnel is 11.43 ± 1.98 for the 1K^+^ simulation and 19.38 ± 2.22 for the 3K^+^ simulation. This suggests that the presence of more K^+^ ions leads to an increase in water molecules in the KdpA portion of the tunnel, most likely because more water molecules are required to hydrate K^+^. The probability distribution of water molecules was also calculated to highlight regions of high water occupancy within the tunnel (Figure 4b). A comparison of the probability peaks from our calculations with the densities observed in the cryo-EM structure is shown in Figure S10. The positions do not precisely overlay because the cryo-EM densities represent individual ordered molecules and the peaks correspond to population densities of mobile molecules. The peaks closely correspond to positions at which densities are observed in the experimental structures. Additionally, the distribution plot clearly shows the hydrophobic region at the interface of the two subunits, where water molecules are absent, and which constitutes the high free energy barrier in the tunnel.

**Figure 4:**
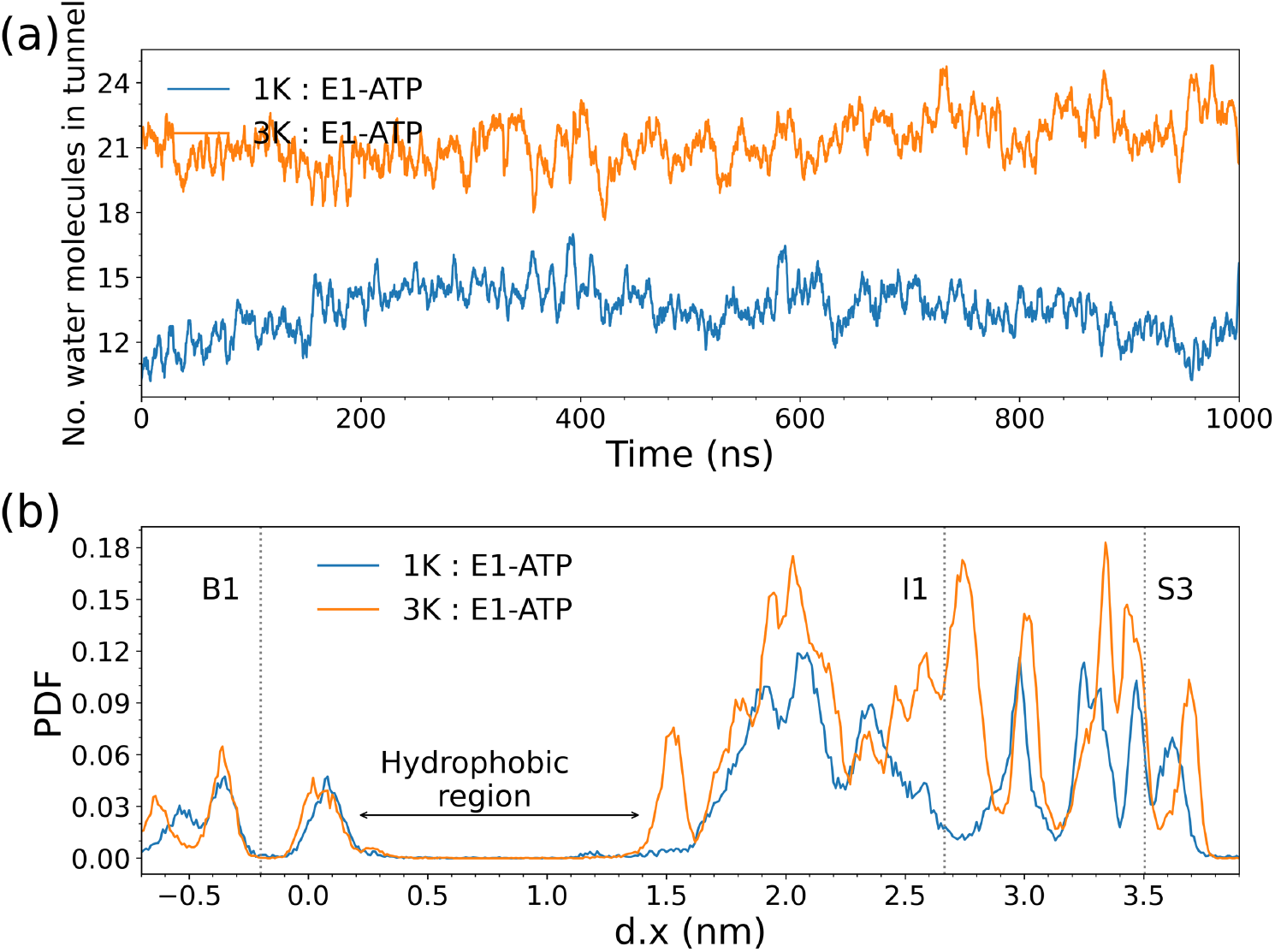
(a) Number of water molecules present in the tunnel when 1 K^+^ or 3 K^+^ ions are initially placed in the tunnel. (b) The probability density of water molecules along the axis of the tunnel. The tunnel region was identified using HOLE.

### *In vitro* activity of mutant KdpFABC

As in previous work by Silberberg et al. ^13^, we have addressed the effects of Phe232 mutation experimentally on transport efficiency and energy coupling. In particular, we compared functional parameters of wild type, F232I, and F232A mutants using assays of ATPase activity and K^+^ transport. Unlike previous work, we used an expression system that prevents complicating effects of post-translational modification, i.e., inhibitory phosphorylation of Ser162 on KdpB, which likely varies depending on the functionality of the construct and the culture conditions^35^. We have compared results from constructs using the WT selectivity filter, which exhibits very high affinity for K^+^ and thus produces background signal in the absence of added K^+^, with the Q116R mutation on KdpA, which lowers the apparent affinity and allows a less ambiguous demonstration of coupling. Finally, we substituted voltage-sensitive fluorescence with solid-supported membrane electrophysiology (SSME) as a more reproducible measure of transport^36, 37^.

Wild type KdpFABC exhibited robust ATPase activity and potassium transport consistent with a well-coupled complex. Although it is not possible to calculate a specific activity based on currents from SSME, typical peak currents of ∼2 nA and ATPase activities of ∼10 μmole/mg/min are consistent with fully coupled pumps under the assumption that proteoliposomes cover ∼25% of the available sensor surface^31^. This coupling is further illustrated by the K^+^ dependence of ATPase activity, which is significantly higher for WT in the presence of 150 mM K^+^. The background activity in the absence of K^+^ is due to the very high affinity of WT (∼5 μM) and K^+^ contamination coming from stock solutions of Mg^2+^ and ATP. To address this problem, we mutated Gln116 in KdpA, which resides at the entrance to the selectivity filter; specifically, the Q116R mutation is well documented to reduce the apparent K^+^ affinity to the millimolar range^9^. For this selectivity filter mutant, there is no ATPase activity in the absence of K^+^, but robust activity in the presence of 150 mM K^+^ (Figure 5). We then measured activities of two F232 mutants, both with the WT selectivity filter and with the Q116R mutation. F232I was used to test the previously proposed cation-pi interaction between K^+^ and the aromatic ring of F232; F232A was used to test the hydrophobic barrier addressed by the simulations. F232I showed strong K^+^-dependence of ATPase activity, suggesting that the pump remained coupled, but a 70-80% reduction in ATPase activity, indicating a reduced efficiency of transport, which is also reflected in a reduced transport current. Consistent with previous analysis, the F232A mutation eliminates the K^+^ dependence of ATPase activity with both WT and Q116R selectivity filters. In addition, transport currents are <5% of WT protein, indicating that the loss of this hydrophobic barrier uncouples the ATP hydrolysis from potassium transport. Unlike the previous analysis, which was susceptible to uncontrolled levels of Ser162 phosphorylation, we do not see a stimulation of ATPase activity, indicating that the hydrolytic domains in KdpB are operating at their maximal rate in the WT construct.

**Figure 5:**
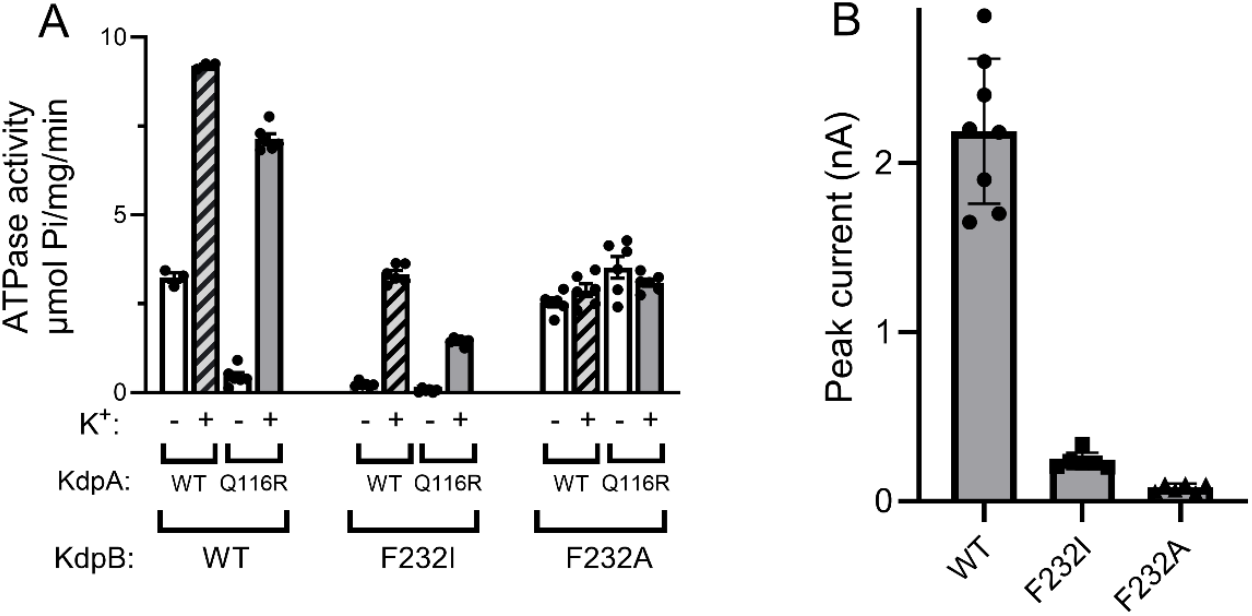
F232A mutant uncouples ATPase from potassium transport. (Left) ATPase activity for wild type (WT) as well as KdpB-F232I and KdpB-F232A mutations. Each of these KdpB mutants was tested with a wild type selectivity filter (KdpA-WT) and a Q116R selectivity filter (KdpA-Q116R) in the presence (+) and absence (−) of 150 mM K^+^. (right) Transport of wild type and the two KdpB-F232 mutants measured with SSME. Uncoupling by the F232A mutation is evident from the K^+^-independent ATPase activity and the complete lack of transport. The F232I mutant retains K^+^-dependence and coupling, but exhibits lower ATPase and transport activities.

### Anomalous X-ray Scattering

Cryo-EM maps have revealed densities in the tunnel, in the selectivity filter and in the canonical binding site in several different conformations.^12, 13^ Although these maps do not distinguish K^+^ ions from water molecules, our simulations indicate that the tunnel is filled with water molecules and that it is energetically unlikely for several K^+^ ions to occupy the tunnel at the same time. To confirm this result, we collected the anomalous signal from X-ray crystallographic experiments to detect K^+^ ions within the structure. Although this approach was previously used to confirm the presence of K^+^ in the selectivity filter of the original X-ray structure ^1^, the 1 Å wavelength used for data collection produced only very weak anomalous signal. We therefore reproduced these crystals and tuned the X-ray radiation to 1.7 Å where this signal is expected to be much stronger. A dataset was collected with diffraction extending to a nominal resolution of 3.1 Å. The resulting anomalous maps (Figure. 6 and Figure. S11) reveal a strong peak at the S3 position of the selectivity filter. The height of this peak ranges from 5-7 sigma for each of the three molecules in the asymmetric unit. This result is a strong confirmation of K^+^ bound to the selectivity filter of KdpA. In the canonical binding site in KdpB, only weak signal (3-4 sigma) is observed, comparable to the noise level of the map; furthermore, the location of the peaks is inconsistent across the three copies in the asymmetric unit and completely absent in one of these molecules, confirming the assignment of water at this site for this inhibited conformation of the pump. In the tunnel, no signal is present above 2 sigma, which is clearly below the noise level of the anomalous map^1^. Therefore, these anomalous X-ray scattering data lend support to the conclusion that the tunnel does not bind multiple K^+^ ions.

**Figure 6:**
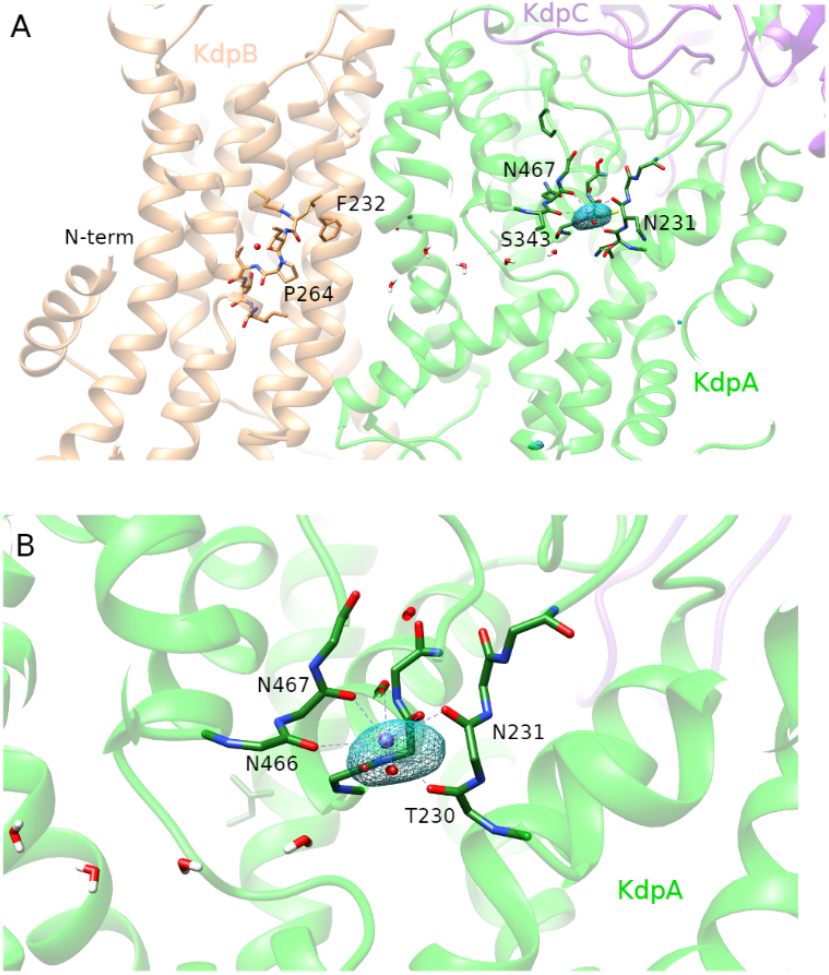
Anomalous signal derived from X-ray crystallography of KdpFABC. The only significant signal was observed in the S3 position of the selectivity filter, shown in closeup in panel B. The mesh surface corresponds to 40 with the peak slightly exceeding 70. No significant signal was seen within the tunnel connecting the subunits. KdpA is colored green, KdpB brown, and KdpC is purple.

## Discussion

Our simulations demonstrate that aside from the selectivity filter sites, only a single intermediate position (I1) can be occupied by K^+^ within the KdpA section of the tunnel. This suggests that most of the densities identified in the tunnel region in cryo-EM studies are likely to be water molecules. The simulations also show that there are not just 7, but between 11-20 water molecules accommodated inside the translocation passage, depending on the number of K^+^ initially placed in the tunnel. We speculate that water molecules which are not tightly coordinated by the protein will not form a well-defined density in the cryo-EM maps, especially considering the limited resolution of those maps (3-3.5 Å). Furthermore, anomalous X-ray scattering maps do not detect any anomalous signal in the tunnel other than a strong signal at S3. Together, our data rule out the existence of a wire of K^+^ ions in the translocation passage. Instead, the densities in the cryo-EM maps correspond to loosely coordinated water molecules. These waters would serve to rehydrate K^+^ after they emerge from the selectivity filter and transiently transit through the tunnel on the way to the canonical ion binding site in KdpB.

Our simulations identified stable binding sites for K^+^ in KdpA subunit at S3, S4 and I1 positions. The S3 site in our study exactly corresponds to the S3 identified by previous cryo-EM studies^12, 13^ and are coordinated by the same residues: N112, T113, T230, S343, C344, N466 and N467. The I1 site detected in our study appears to correspond to the T2 site identified by Silberberg et al^13^ and possibly corresponds to either A3 or A5 site labelled by Sweet et al^12^, which are coordinated by the same residues in the PDB IDs 7NNL and 7LC3, respectively. Although this appears as a relatively stable site in our simulations, the energy profile is relatively flat in this region, consistent with this site as a transient waypoint as the ion travels through the tunnel. The S4 site in our study was coordinated by residues A342, S3433, G374, S378 and N466. The S4 site ascertained by Silberberg et al^13^ is also coordinated by S343, S378 and N466. Thus, our findings reveal a new site I1 and corroborate the previously identified S3 and S4 sites. In our simulations, when only two K^+^ ions were present in the tunnel, they always occupied S3 and I1. Therefore, it can be concluded that S4 is the least stable binding site among the three sites. The B site in the KdpB subunit is analogous to the CBS identified by Silberberg et al^13^ and may possibly be Bx or B1 identified by Sweet et al^12^.

Our simulations failed to detect K^+^ translocation through the tunnel, which can be explained by the presence of a large free energy barrier near the interface of KdpA and KdpB subunits. Metadynamics simulations indicate this barrier to be >22 kcal/mol, which is almost 20-fold higher than thermal energy available to the system (*k*T). This result contrasts with previous simulations reported by Silberberg and co-workers^13^ in which translocation across the interface readily occurred. There are four key differences between the two simulation systems. First, the initial simulation setup in Silberberg et al.^13^ included a large number of K^+^ ions in the tunnel, which would be expected to generate considerable electrostatic repulsion. Given that the energies reported by our metadynamics are relative, a high initial energy would make it possible to overcome the barrier at the hydrophobic interface. Thus, this electrostatic repulsion is likely to account for the observed transfer of K^+^ across the interface, which occurred on a very short timescale (few picoseconds). Furthermore, the simulations were too short to observe escape of K^+^ from any other route such as the selectivity filter as in our simulations. We contend that a state with several K^+^ ions in the translocation passage is thermodynamically unfavorable and thus not consistent with physiological conditions. Secondly, in Silberberg’s simulations, Lys586 near the KdpB binding site was kept uncharged. In our simulations we have kept Lys586 charged according to the pKa value of the residue as calculated using PropKa. In our simulations, a positive charge on Lys586 may resist the binding of K^+^ to the KdpB binding site. We and others have previously shown that the ionization states of the residues at the ion-binding site in P-type ATPases can change depending on the conformational state in the Post-Albers cycle ^33, 38, 39^; therefore, conformation-dependent dynamic ionization state of Lys586 cannot be ruled out. Thirdly, in some simulations of Silberberg et. al^13^, the cytoplasmic domains of protein were truncated, and the protein backbone atoms were restrained, which would prevent any local structural changes in the protein which may occur in response to the unfavorable placement of K^+^ ions. In contrast, we have simulated the entire protein without any position restraints for all simulations. Finally, free energy profiles calculated using Metadynamics are significantly more accurate than the semi-empirical APBS method used by Silberberg et. al.^13^

Our metadynamics simulations indicate a surprisingly high barrier for the transfer of a K^+^ ion across the hydrophobic interface: ∼ 22 kcal/mol. We observed the high barrier for ion translocation along the interface for each of the E1-ATP and E1-ATP-F232A states. The free energy of ATP hydrolysis under cellular conditions (∼ 14 kcal/mol) cannot overcome this barrier. The high barrier in our simulations must be interpreted in two contexts: first, the energy profile is calculated along a CV which resembles the predicted movement through the tunnel but may not exactly conform to the lowest energy path. It is possible that movement in a different direction would reduce the magnitude of the barrier. Secondly, the lack of water molecules or hydrophilic protein ligands at the interface implies that transfer of a K^+^ would require at least partial dehydration, which involves a very high free energy cost. The theoretical hydration free energy of K^+^, which quantifies the free energy cost of removing a K^+^ from water and placing it in vacuum is ∼ 70-75 kcal/mol^40^. To facilitate transport across the barrier, the K^+^ ion must then be coordinated temporarily with a polar molecule such as an amino acid side chain or water molecule(s) at the hydrophobic interface, such as the carbonyl oxygens that characterize the selectivity filter. We speculate that there is an intermediate state of the KdpFABC between the known configurations which lowers the barrier for ion transport by providing such a polar species. Such a state, which may be too transient to be seen under turnover conditions and not stabilized by the inhibitors used in previous structural work, remains to be identified.

In conclusion, our investigations show that the tunnel extending from KdpA to KdpB in KdpFABC is filled not with a queue of K^+^ ions, but with a considerable number of water molecules. Densities in the cryo-EM maps of previously solved structures correspond to a few of the better ordered water molecules at sites that provide coordination from the surrounding protein moieties. In the E1-ATP state, the free energy barrier of transport of K^+^ along the tunnel is too high for spontaneous ion transfer to occur. The exact mechanism by which K^+^ switches from the KdpA-KdpB interface to the KdpB canonical ion binding site remains to be identified.

## Acknowledgements

HVM and HK are supported by the Lundbeckfonden Ascending Investigator grant (#R344-2020-1023). The MD simulations were carried out on the Danish e-Infrastructure Cooperation (DeiC) National HPC Center, on ABACUS 2.0 at the University of Southern Denmark, Novo Nordisk Foundation funded ROBUST Resource for Biomolecular simulations #NNF18OC0032608, finnish supercomputer LUMI, under grant number DeiC-SDU-N5-000007, and Vega supercomputer, Slovenia under the EUROHPC grants #EHPC-REG-2021R0060 and #EHPC-REG-2022R02-207

DLS is supported by the National Institutes of Health (Grant Agreement No. R35 GM144109).

The authors acknowledge beamlines I24 and I04 at the Diamond Light Source, beamline BioMAX at the MAX IV Laboratory and DESY-PETRA III for crystal screening. We acknowledge the MAX IV Laboratory for beamtime on the BioMax beamline under proposal 20200251 for final data collection. Research conducted at MAX IV, a Swedish national user facility, is supported by Vetenskapsrådet (Swedish Research Council, VR) under contract 2018-07152, Vinnova (Swedish Governmental Agency for Innovation Systems) under contract 2018-04969 and Formas under contract 2019-02496.

