## Supplemental figures and tables for "On the mechanism of K^+^ transport through the inter-subunit tunnel of KdpFABC"

**[2] Dept. of Molecular Biology and Genetics, Aarhus University, 8000 Aarhus C, Denmark**

**[3] Dept. of Biochemistry and Molecular Pharmacology, New York University School of Medicine, New York, NY 10016 USA**

**Table 1:** Data collection and refinement statistics

| | |
|---|---|
| **Name** | KdpFABC Q116R (xtal data is DS25-D1e) |
| **State** | E1-P |
| **Data Collection** | |
| Space group | P 21 |
| Cell dimensions | |
| a. b, c (Å) | 121.0, 165.2, 192.6 |
| alpha, beta, gamma (deg) | 90 105.5 90 |
| Monomers per asym. unit. | 3 |
| Wavelength (Å) | 1.7712 |
| Number of reflections measured | 381300 |
| Number of unique reflections | 55764 |
| Resolution (Å) | 185.6-3.1 (3.4-3.1)a |
| Rmeas (%) | 11.2 (101.0) |
| Mean I/σ(I) | 10.5 (2.1) |
| CC(1/2) | 99.5 (63.9) |
| Completeness (spherical) (%) | 41.5 (7.5) |
| Completeness (ellipsoidal) (%) | 80.6 (74.7) |
| Redundancy | 6.8 (7.0) |

[a] Highest resolution shell is shown in parenthesis.

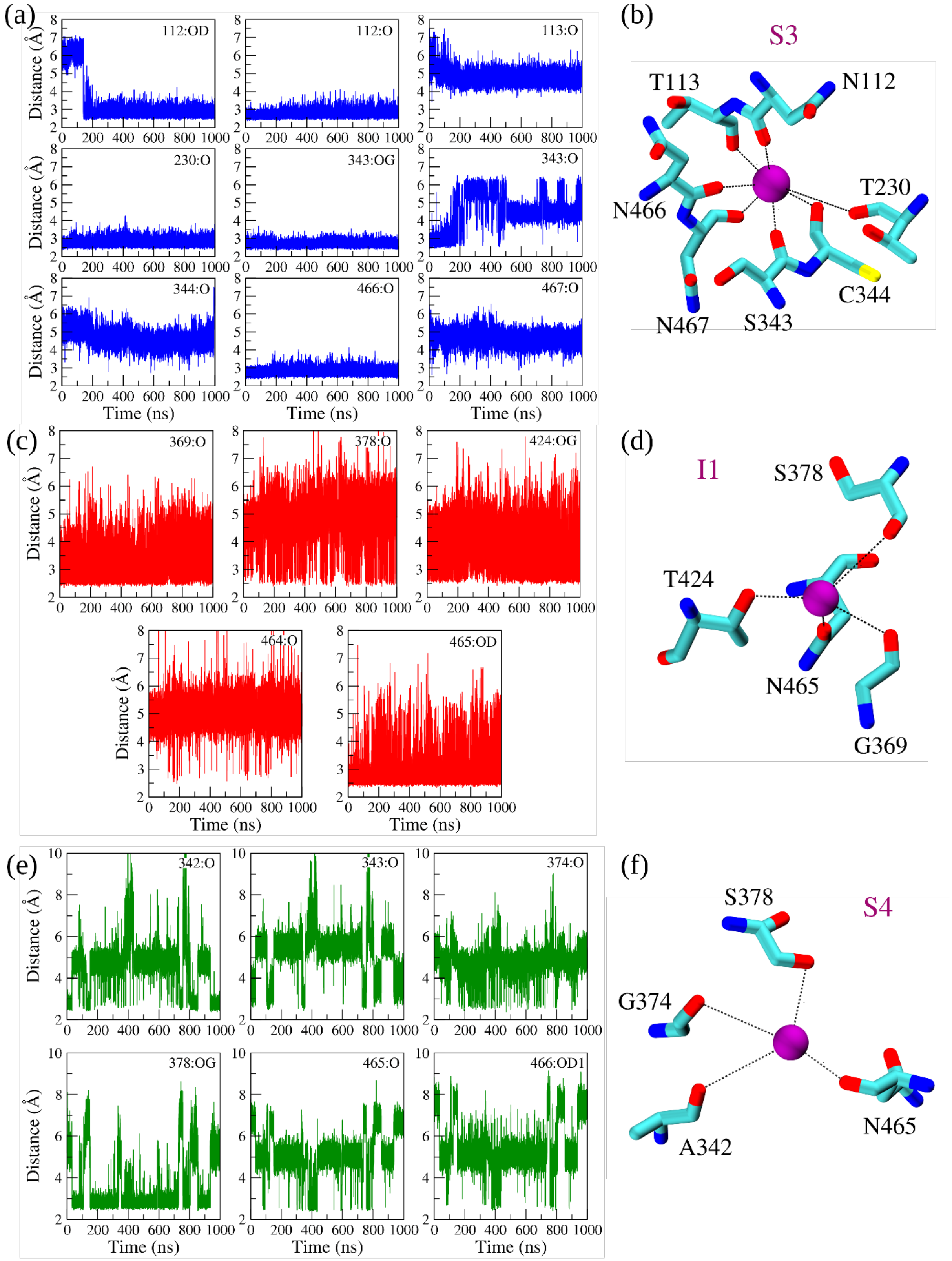

**Figure S1**: The distance of the $K^+$ ion from the atoms of the residues that coordinate the $K^+$ ion at different sites in the tunnel (a) S3, (c) I1 and (e) S4 along the simulation time. Panels (b), (d) and (f) shows the coordination of the residues to the $K^+$ ion.

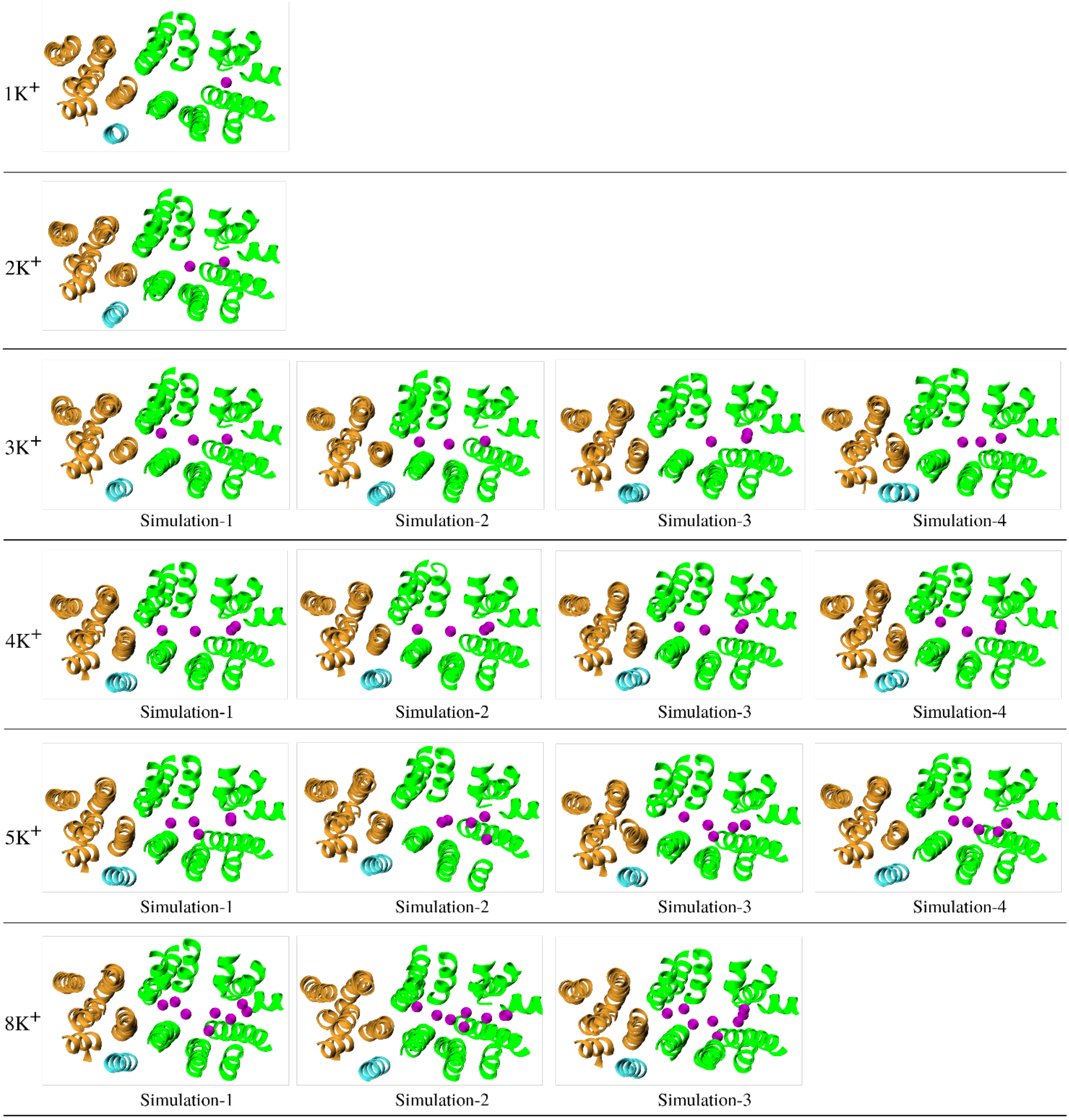

**Figure S2**: Positions of the $K^+$ ions in the tunnel at the beginning of the simulation in the different simulation sets.

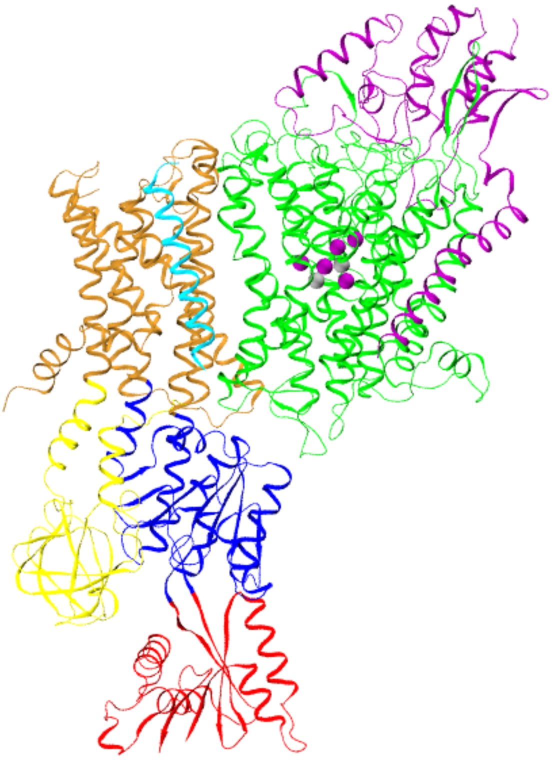

**Figure S3**: Presence of more than 3 $K^+$ ions in the KdpA part of the translocation passage attracts $Cl^-$ ions from the solvent. The $K^+$ ions are shown in purple and the $Cl^-$ ions are shown in silver.

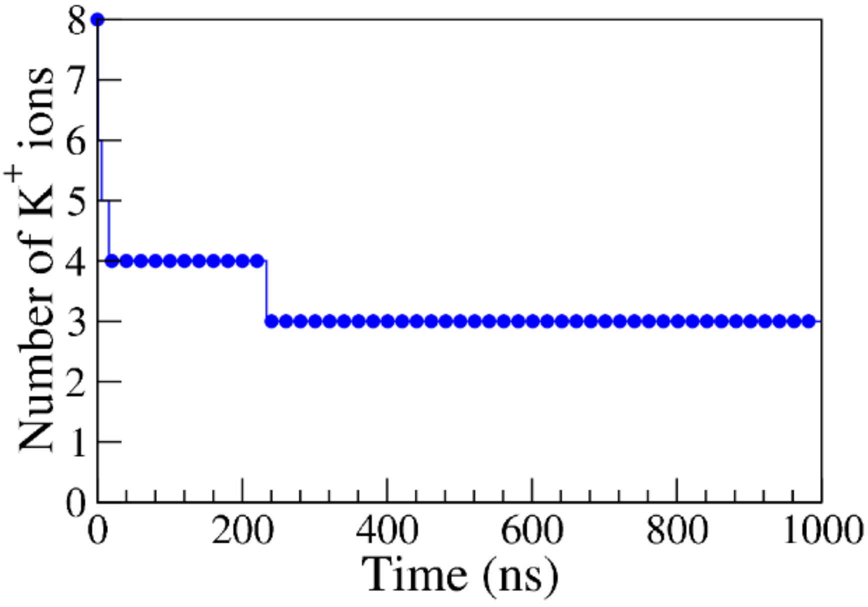

**Figure S4**: Number of $K^+$ ions that remain in the translocation passage in KdpFABC-F232A simulation.

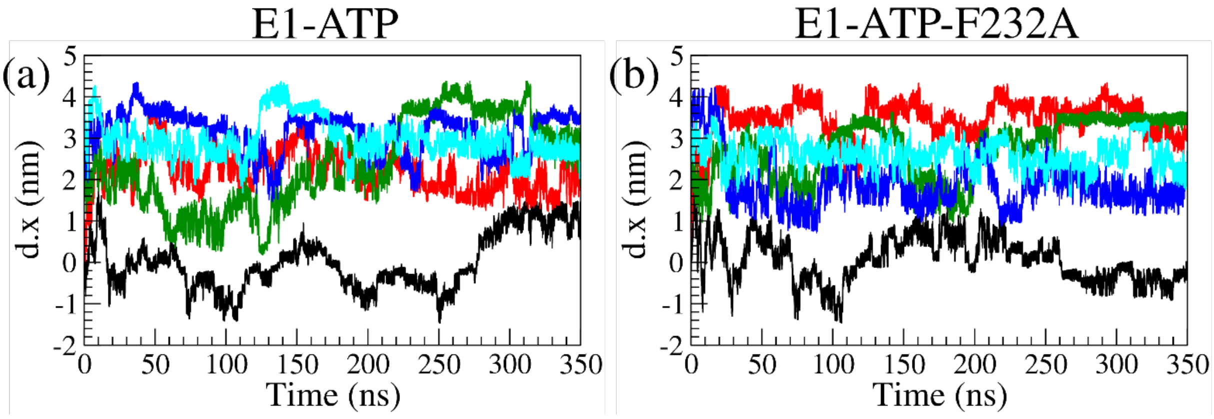

**Figure S5**: Evolution of the reaction coordinate d.x (projection of distance of $K^+$ ion from B site along the x-axis in nm) as a function of simulation time for the five different walkers in the metadynamics simulations of the protein.

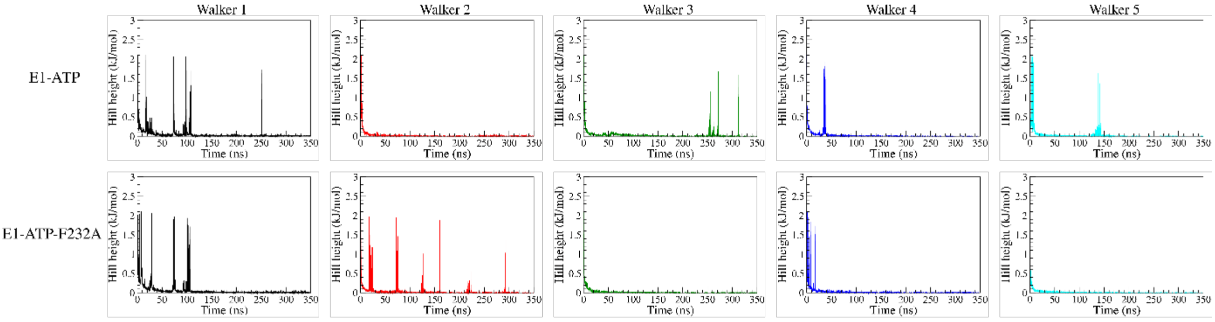

**Figure S6**: Hill height along the simulation time for all the five walkers in the two systems. The significant hill depositions after 100 ns in most of the walkers correspond to sampling in the region of disinterest (near the walls).

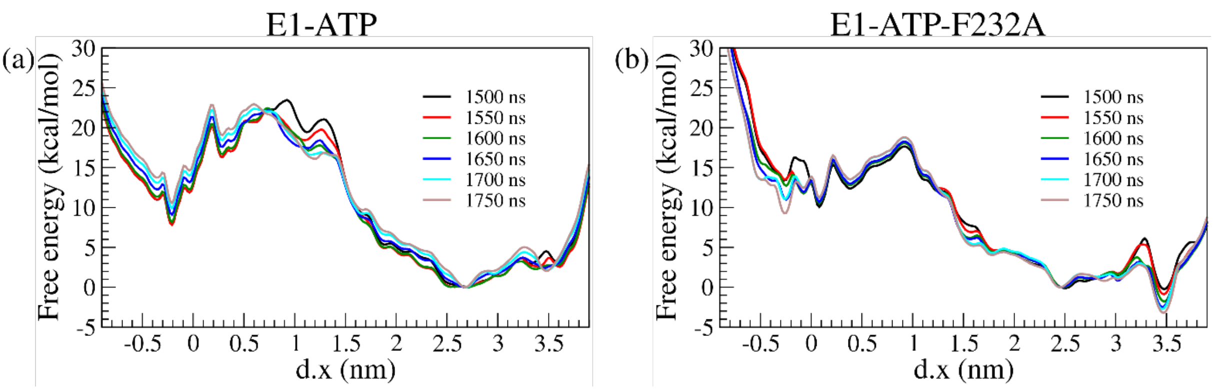

**Figure S7**: Convergence of the free energy in the systems studied.

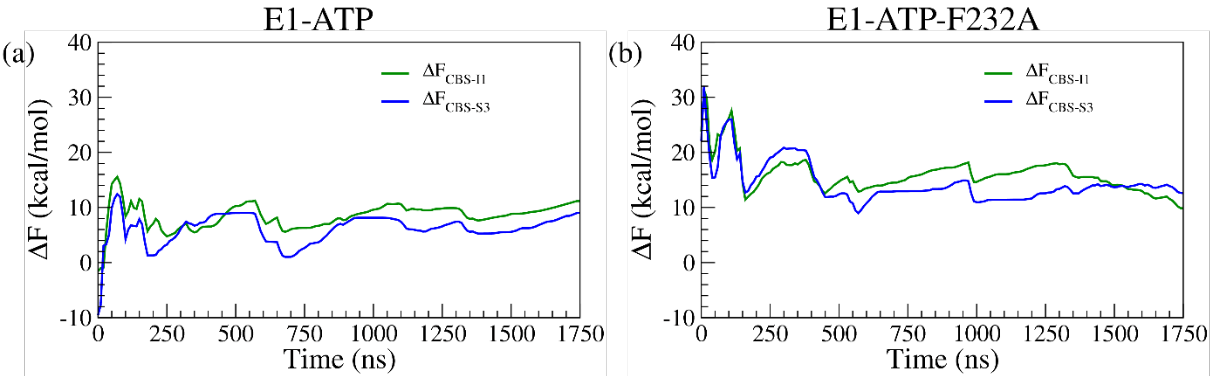

**Figure S8**: Free energy convergence: Evolution of free energy differences, $\Delta F_{CBS-I1}$ and $\Delta F_{CBS-S3}$ as a function of simulation time.

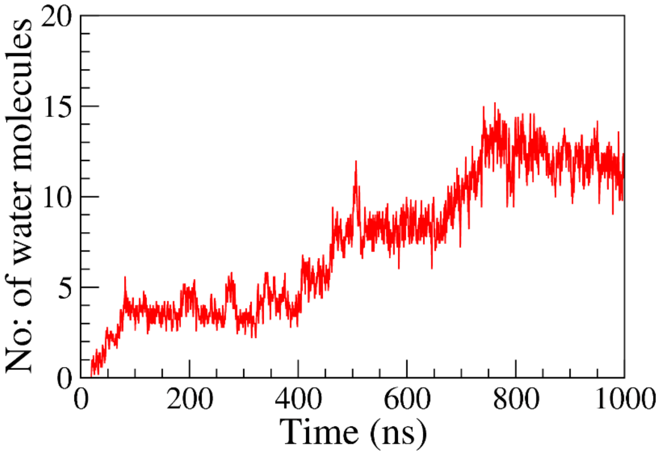

**Figure S9**: The number of water molecules in the tunnel of E1-ATP conformation of the protein as a function of the simulation time. The figure depicts a running average of every 5 data points in the simulation. The tunnel boundary was determined using HOLE.

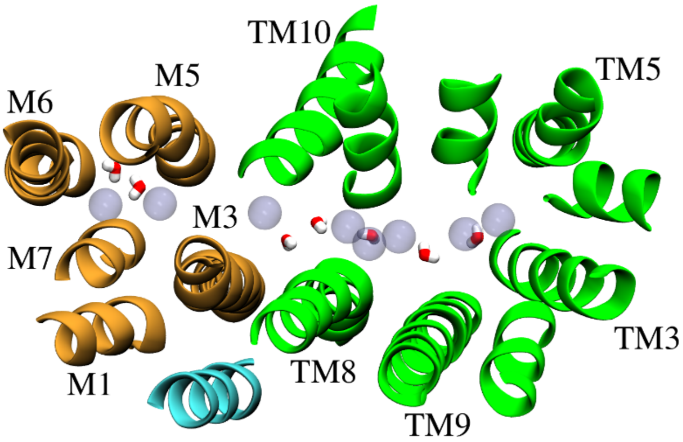

**Figure S10**: Comparison of water densities in the tunnel: the water molecules present in the cryo-EM (PDB ID: 7LC3) structure is represented by licorice model whereas the positions where water molecules were highly likely to be found in the simulation (corresponding to the peaks in Figure 4b in manuscript) are represented by spheres.

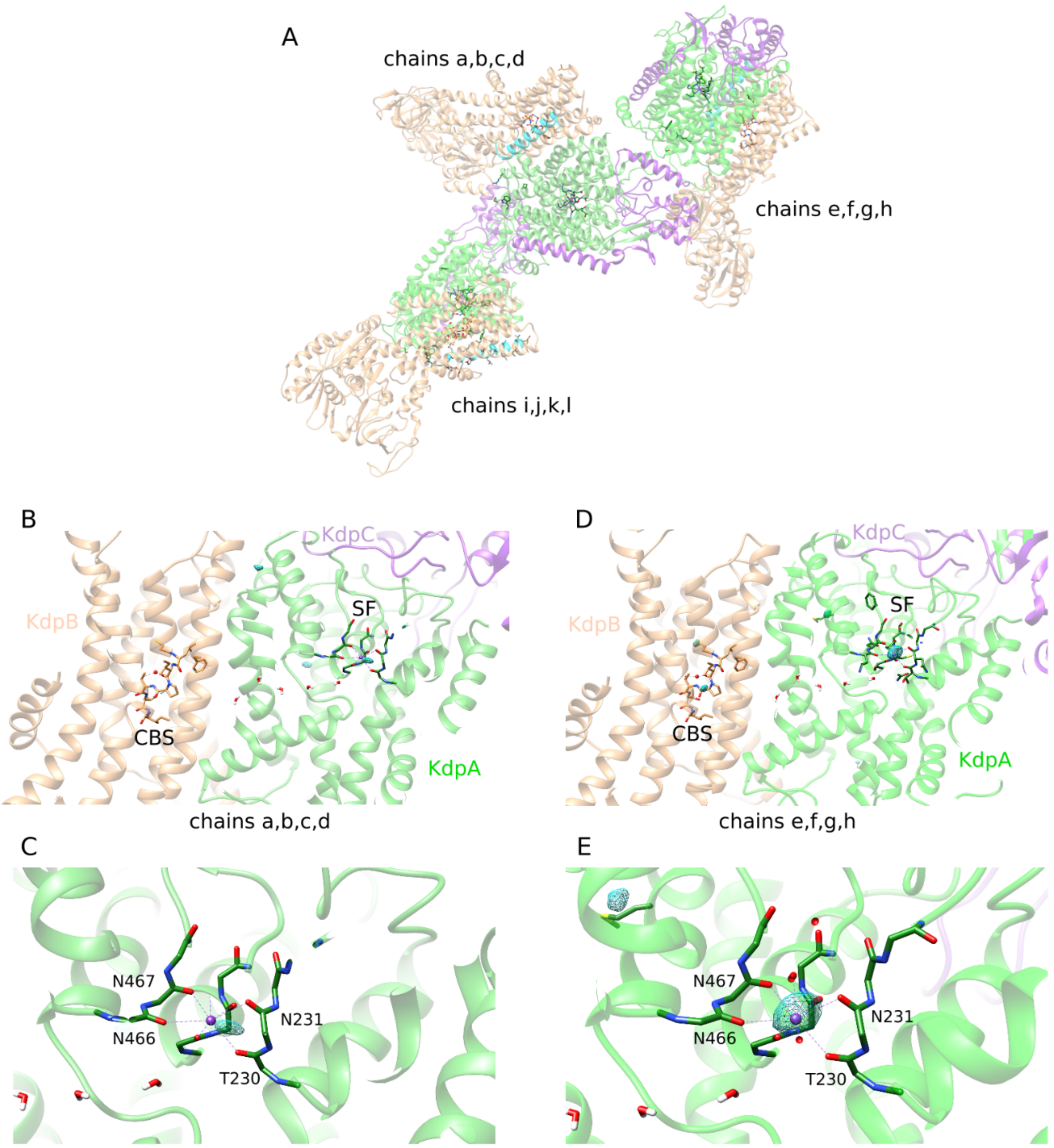

**Figure S11**: Anomalous signal produced by X-ray diffraction of KdpFABC crystals. (A) Three copies of KdpFABC compose the asymmetric unit. Data for chains i,j,k,l are shown in Fig. 6 whereas data for the other two, independent molecules are shown below. (B,C) Anomalous signal for chains a,b,c,d is shown by the mesh surface representing 4s. The peak maximum at S3 is 4.9. (D,E) Anomalous signal for chains e,f,g,h is shown by the mesh surface representing 4s and maximal signal of 6.1s. The ribbons are colored green for KdpA, brown for KdpB, purple for KdpC, and cyan for KdpF.
